# GuiltyTargets: Prioritization of Novel Therapeutic Targets with Deep Network Representation Learning

**DOI:** 10.1101/521161

**Authors:** Özlem Muslu, Charles Tapley Hoyt, Martin Hofmann-Apitius, Holger Fröhlich

## Abstract

The majority of clinical trial failures are caused by low efficacy of investigated drugs, often due to a poor choice of target protein. Computational prioritization approaches aim to support target selection by ranking candidate targets in the context of a given disease. We propose a novel target prioritization approach, GuiltyTargets, which relies on deep network representation learning of a genome-wide protein-protein interaction network annotated with disease-specific differential gene expression and uses positive-unlabeled machine learning for candidate ranking. We evaluated our approach on six diseases of different types (cancer, metabolic, neurodegenerative) within a 10 times repeated 5-fold stratified cross-validation and achieved AUROC values between 0.92 - 0.94, significantly outperforming a previous approach, which relies on manually engineered topological features. Moreover, we showed that GuiltyTargets allows for target repositioning across related disease areas. Applying GuiltyTargets to Alzheimer’s disease resulted into a number of highly ranked candidates that are currently discussed as targets in the literature. Interestingly, one (COMT) is also the target of an approved drug (Tolcapone) for Parkinson’s disease, highlighting the potential for target repositioning of our method.

**Availability:** The GuiltyTargets Python package is available on PyPI and all code used for analysis can be found under the MIT License at https://github.com/GuiltyTargets.

**Author summary:** Many drug candidates fail in clinical trials due to low efficacy. One of the reasons is the choice of the wrong target protein, i.e. perturbation of the protein does not effectively modulate the disease phenotype on a molecular level. In consequence many patients do not demonstrate a clear response to the drug candidate. Traditionally, targets are selected based on evidence from the literature and follow-up experiments. However, this process is very labor intensive and often biased by subjective choices. Computational tools could help a more rational and unbiased choice of target proteins and thus increase the chance of drug discovery programs. In this work we propose a novel machine learning based method for target candidate ranking. The method (GuiltyTargets) captures properties of known targets to learn a ranking of candidates. GuiltyTargets compares favorably against existing machine learning based target prioritization methods and allowed us to propose novel targets for Alzheimer’s disease.

## Introduction

Drug discovery is a time consuming, expensive and complicated process [1–4]. Many drug candidates fail in clinical studies due to low efficacy, mainly because of wrong target choice [5–7]. Traditionally, scientists identified targets by searching through the relevant literature, following clues from mRNA and protein expression, integrating expression data with pathway analyses, experimenting with knockout mice, investigating somatic mutations, gene fusions, and copy number variations, and using the accumulated knowledge from multiple experimental studies to generate a hypothesis on how a molecule might work as a target [2, 8, 9]. However, manually interpreting many data sources is prone to biased identification of targets as it limits the potential to use all available and helpful data. By computationally integrating multiple biological data sources to analyze prior knowledge, it should be possible to make target identification process faster, less biased, and more informed. Computational target prioritization approaches thus aim for improving target identification process by ranking proteins based on their likelihood of being targets in the context of a specific disease [10–17]. Most of them integrate biological networks with other data sources to prioritize targets for infectious diseases [11–13], cancers [14–16] or neurodegenerative diseases [17].

In addition, machine learning methods have been used to prioritize drug targets. For example, Emig *et al*. proposed an approach, in which for each candidate target a number of different network topological features are combined with proximity to differentially expressed genes in a particular disease of interest [10]. All features are subsequently combined into a logistic regression model, which allows for a ranking of candidate targets. The authors successfully tested their approach with 30 different diseases. Another example is the method by Ferrero *et al*., which uses features provided by the Open Targets database [18] and combines them into one ranking score using Support Vector Machines [19].

In this paper we propose a novel approach to prioritize targets using a combination of unsupervised network representation learning, namely the recently proposed Gat2Vec method [20], and logistic regression. More specifically, our method, GuiltyTargets, first maps a genome-wide protein-protein interaction network annotated with differential gene expression information into an Euclidean space using Gat2Vec. In that space, we then use positive-unlabeled (PU) machine learning [21–24] to learn a ranking of candidate targets. To the best of our knowledge, network representation learning as a data driven approach to implicitly learn relevant topological features from a network structure has not been used for target prioritization so far. The proposed approach is compared to the approach from Emig *et al*. [10] for six diseases, demonstrating its superior ranking performance. For the example of Alzheimer’s disease (AD), in-depth analysis shows that GuiltyTargets can be used to reposition known targets from other neurological indications.

## Results

### GuiltyTargets: A New Approach for Deep Network Representation Learning Based Target Prioritization

Our newly developed GuiltyTargets method can be summarized as follows (Figure 1): First, a genome-wide PPI network is compiled and annotated with discretized information about differential gene expression within a given disease context (-1 = underexpressed; 0 = no significant change; 1 = overexpressed). Next, the attributed network is embedded into an Euclidean space using Gat2Vec. Following a PU learning scheme known disease specific protein targets are assigned positive labels, and the remaining proteins are regarded as pseudo-negatives to train a classifier that ranks a candidate protein according its similarity to known targets for the given disease. More details about GuiltyTargets are described in the Methods section of this paper.

**Fig 1.**
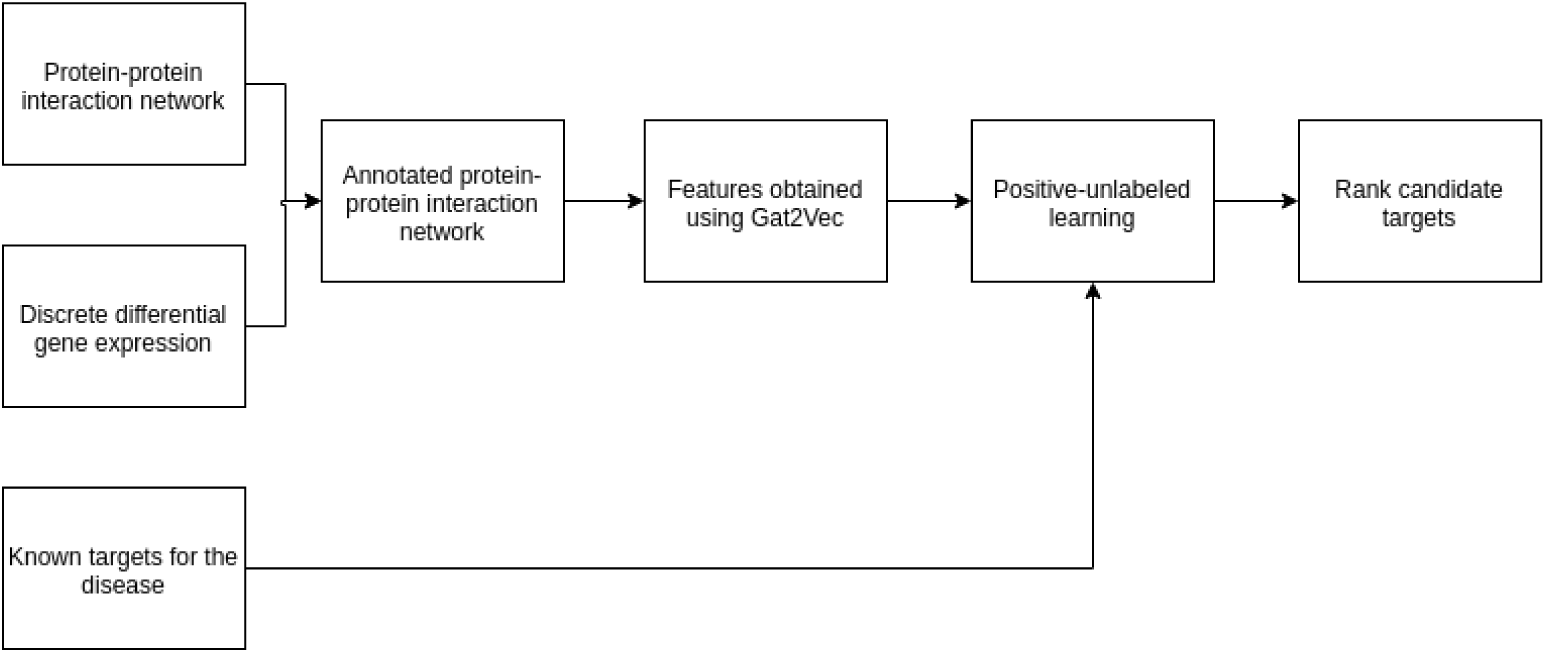
GuiltyTargets pipeline. First, a protein-protein interaction network is annotated with discrete differential gene expression information (up-regulated, down-regulated, not differentially expressed). 128 features are extracted from this annotated network using Gat2Vec. These features are input to a logistic regression algorithm where known targets are labeled as positive and the remaining as negative. The likelihoods that are calculated by the classifier is then used for ranking.

### Validation Data

We performed target prioritization analyses for six different diseases using corresponding gene expression data for acute myeloid leukemia, hepatocellular carcinoma, idiopathic pulmonary fibrosis, liver cirrhosis, multiple sclerosis and AD. The choice was made based on the following criteria: First, five of these diseases have also been evaluated in the publication by Emig *et al*. [10], which we used for comparison here. Second, the number of available known targets for each disease was expected to be relatively high for a statistically meaningful validation. Finally, we added AD to investigate the applicability of our approach to a highly challenging disease, in which so far most attempts to establish new drugs have failed [25]. Details about about used data, including pre-processing, are described in Materials and Methods Section. Notably, for AD we investigated RNASeq data from different cohorts (MSBB [26], MayoRNASeq [27], ROSMAP [28]). To investigate the prediction performance of GuiltyTargets we employed two protein-protein interaction networks (STRING [29], HIPPIE [30]), two target databases (Open Targets [18], Therapeutic Target Database [31]) and different cutoffs to discretize differential gene expression via *log*_*2*_ fold change thresholds (0.5, 1.0, 1.5) while requiring a false discovery rate of less than 5%.

### GuiltyTargets Outperforms Existing Method

The performance of our approach and the method by Emig *et al*. were compared within a 10 times repeated 5-fold cross validation scheme with the area under ROC curve (AUROC) as the evaluation criterion. This assessed the ability of each method to rank in an independent test set a true known target higher than an unknown protein. For this purpose, the approach employed by Emig *et al*. was re-implemented using the same PPI network resources and target databases as for GuiltyTargets.

Results shown in Tables 1, S1 and S2 demonstrate a dramatic performance increase of up to 40% by GuiltyTargets compared to the method by Emig *et al*.. Notably, AUROC values found by our re-implementation of were not identical (but typically close) to the ones reported in the original paper. This was likely due to the fact that not the same PPI network and target database resources have been used. More specifically, Emig *et al*. employed the commercial MetaBase™database, whereas here we only rely on public resources.

**Table 1.** Ranking performance of GuiltyTargets compared to the method by Emig *et al*. in terms of cross-validated AUROC (standard error). The table shows the results when using Open Targets as resource for drug targets, STRING (using the default confidence threshold) as PPI network and declaring differential gene expression based on log2 fold change cutoff of 1.5, which is agreement to Emig *et al*. The column “Reported” shows the AUROC values reported by Emig *et al*. and the column “Implemented” shows the AUROC values obtained by the reimplementation of their method. The row “Alzheimer’s Disease” refers to the MayoRNAseq data from temporal cortex, which among all tested AD gene expression datasets showed genes with an absolute log2 fold change larger than 1.5.

| Disease | Emig *et al.* (original) | Emig *et al.* | GuiltyTargets |
| --- | --- | --- | --- |
| Acute myeloid leukemia | 0.8195 | $0.8356 \pm 0.0001$ | $0.9277 \pm 0.0002$ |
| Alzheimer’s disease | - | $0.6235 \pm 0.0010$ | $0.9418 \pm 0.0004$ |
| Hepatocellular carcinoma | 0.8019 | $0.7314 \pm 0.0002$ | $0.9384 \pm 0.0001$ |
| Idiopathic pulmonary fibrosis | 0.8826 | $0.8306 \pm 0.0018$ | $0.9263 \pm 0.0004$ |
| Liver cirrhosis | 0.6747 | $0.5338 \pm 0.0019$ | $0.9464 \pm 0.0002$ |
| Multiple sclerosis | 0.7151 | $0.6755 \pm 0.0002$ | $0.9412 \pm 0.0003$ |

We employed an ANOVA to assess the statistical significances of our findings. The ANOVA model was built separately for each investigated scenario (PPI network, target database, log2 fold change cutoff) with three factors: method, dataset and an interaction term between method and dataset. The ANOVA F-test confirmed a highly significant improvement of GuiltyTargets compared to the method by Emig *et al*. for each disease and scenario (*p* < 0.001 after Holm’s correction for multiple testing, see Figure 2). As expected there was a highly significant dataset dependency of AUROC performance in every case (*p* < *-*2.2*e* 16). Post-hoc analysis using Tukey’s multiple comparisons of means revealed that, on average, GuiltyTargets outperformed the approach by Emig *et al*. by 14.8% AUROC.

**Fig 2.**
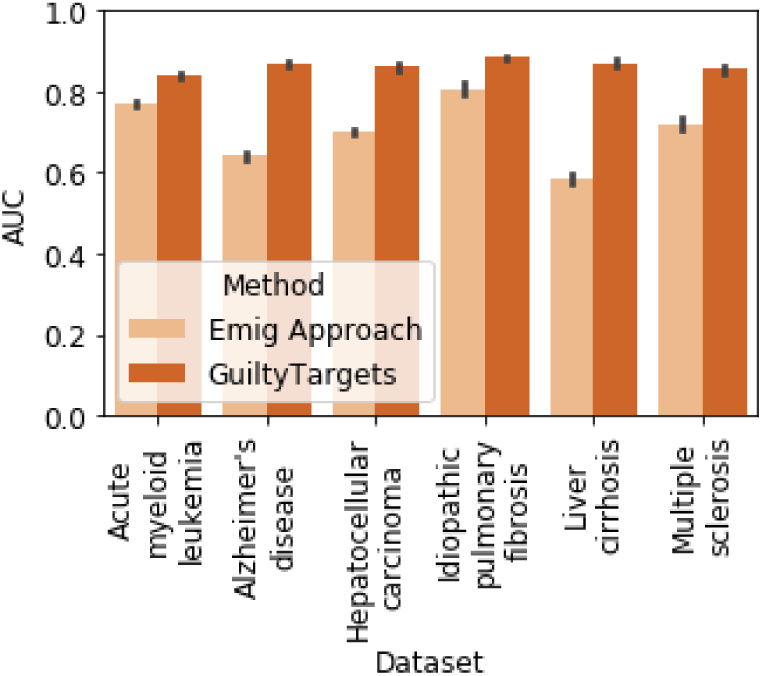
frohlich: Please change legend: Comparison of GuiltyTargets vs approach by Emig *et al*.: The barplots show AUC values averaged over all possible hyper-parameter choices (log2 fold change cutoff, target database, PPI network, PPI network confidence cutoff).

### In-Depth Analysis of Influence Factors on GuiltyTargets Performance

We wanted to better understand the dependency of the performance of GuiltyTargets on the different tested influence factors, which we had varied individually in our cross-validation analysis:

- PPI network (STRING, HIPPIE), including different confidence level thresholds
- Target database (Open Targets, Therapeutic Targets Database)
- Thresholds for declaring differential gene expression

For this purpose we fitted a two-way ANOVA model with interaction term (influence factor, dataset, interaction term between both) and then performed a Tukey post-hoc analysis. Table 2 demonstrates that the employed PPI network is the most relevant influence factor for GuiltyTargets: Using STRING significantly increased the AUROC compared to using HIPPIE by 9%. On the other hand, the chosen log2 fold change threshold had almost no influence, i.e. GuiltyTargets is highly robust against this parameter. A more conservative confidence threshold for the STRING network yielded a drop in prediction performance by 3%. Both findings together may be explained by a comparably strong influence of the network topology for GuiltyTargets, which is leveraged by Gat2Vec. An obvious question is therefore, in how far GuiltyTargets is affected at all by gene expression data or whether our method is purely topology based. We thus compared the performance of our method across the three tested AD gene expression datasets (MSBB, MayoRNASeq, ROSMAP), confirming a significant influence of the actually used dataset on the AUROC for this particular disease (*p* = 3.09*E -* 09, ANOVA F-test). Hence, gene expression data does have a clear effect on GuiltyTargets.

**Table 2.** In-depth analysis of different influence factors on the performance of GuiltyTargets. The table shows the result of the Tukey ANOVA post-hoc analysis of different comparisons. The difference in AUROC is shown in column 3 together with the corresponding p-value in column 4.

| Influence factor | Comparison | Difference | p-value |
| --- | --- | --- | --- |
| PPI network | STRING vs. HIPPIE | 0.09 | 1.15E-14 |
| PPI confidence threshold | default vs 0.63 | 0.0318 | 1.15E-14 |
| Target database | Open Targets vs. TTD | 0.0527 | 2.23E-10 |
|  | 0.5 vs 1.0 | -0.0016 | 0.01369 |
| Log2 fold change cutoff | 0.5 vs 1.5 | 0.0007 | 0.46799 |
|  | 1.0 vs 1.5 | 0.0023 | 0.00026 |

The use of the Open Targets versus the Therapeutic Target Database significantly increased the ranking performance of GuiltyTargets by 5%, hence underlining the relevance of a larger number of known targets for learning the ranking model in the embedded network space.

### GuiltyTargets Learns from Known Targets

We tested, whether the good performance of GuiltyTargets was dependent on known targets or whether also with a random set of proteins a similar performance could have been achieved. For this purpose we trained GuiltyTargets for each disease with 100 randomly drawn sets of targets of the same size as the actual ones, which we incorrectly labeled as “targets”. Prediction performance was evaluated using the same cross-validation procedure as before. Table S3 confirms that the AUROC for random proteins drops to about 50%, i.e. chance level. Hence, GuiltyTargets indeed learns properties of known targets.

### GuiltyTargets Allows for Target Repositioning Across Related Diseases

There is the question whether GuiltyTargets could transfer properties learned from known targets in one disease area into another one, hence allowing for repositioning of targets. To address that question we trained GuiltyTargets with all known targets of neurodegenerative diseases obtained from Open Targets, while excluding known AD targets. We then ran a hypergeometric test on the resulting prioritization to see if known AD targets where statistically overrepresented at the top of the list. The results were significant when at least 2% of the top candidates were considered (Figure 3). This shows that GuiltyTargets could help for repositioning targets across related disease areas.

**Fig 3.**
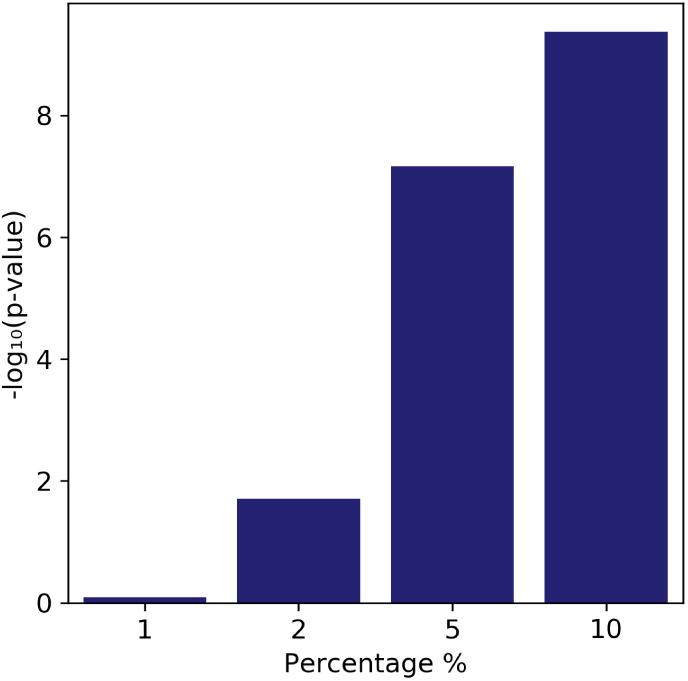
Target repositioning potential of GuiltyTargets: The barplot shows the result of a hypergeometric test conducted on the top p% of a ranked list of candidate proteins when looking for overrepresentation of known AD targets. GuiltyTargets was trained without any known AD targets here.

### Case Study: GuiltyTargets Predicts New Candidate Targets for Alzheimer’s Disease

Despite of 180 therapeutic targets listed in the Open Targets database, the AD field urgently requires new and more effective medications that either prevent, mitigate, or reverse its progression. This is particularly true, because the vast majority of drugs under development fail in clinical trials [25]. We here picked out AD as a test case for GuiltyTargets to prioritize new target candidates. We used post-mortem gene expression data from brain tissue from the ROSMAP study and combined it with STRING based PPI network and Open Targets as a resource for known targets. ROSMAP data was chosen because of its comparably large number of samples (495 AD patients and 438 controls). Table 3 shows the top 0.1% of a ranked list of novel candidate targets (likelihood score > 0.4) obtained with GuiltyTargets. According to the TTD [31] and DGIdb [32] databases, all but two candidates are druggable, thus they could be used as targets for drugs using the current drug development methods.

**Table 3.**
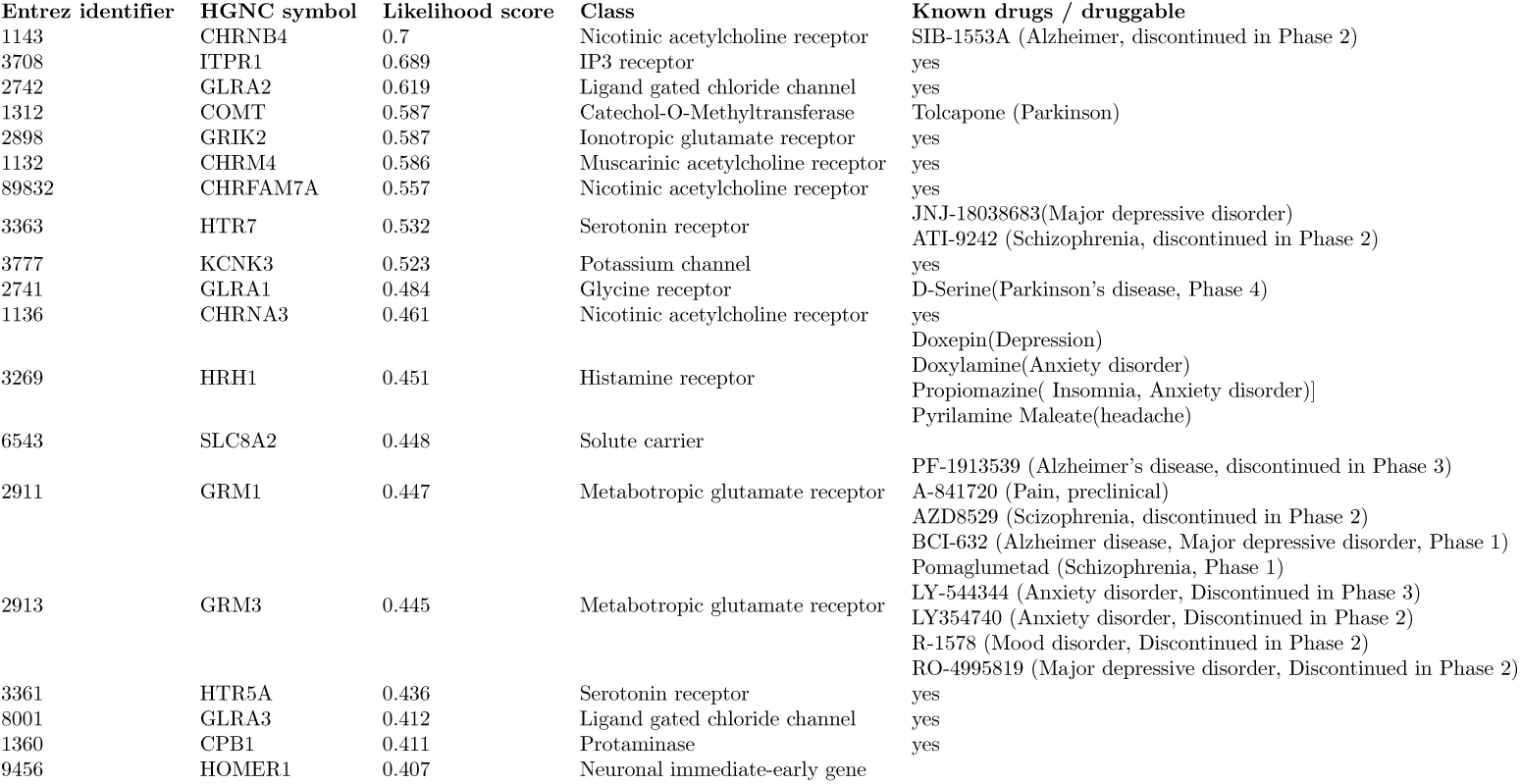
Target prioritization for AD using ROSMAP gene expression data. This list shows candidate proteins above a likelihood score threshold of 0.4. The last column shows either known drugs (including indications) against the respective target or the classification as “druggable” using the information from DGIdb [32] and TTD [31].

Many of the candidate targets are receptors, namely four acetylcholine receptors (three nicotinic, one muscarinic), and three glutamate receptors (two metabotropic, one ionotropic), in agreement with the observation that receptors constitute a large portion of known AD targets for small molecule drugs [33]. The remaining candidates were identified as ion channels. The top candidate (CHRNB4) is the target of the compound SIB-1553A, which has been tested in a phase 2 clinical trial for AD, but discontinued (source: Therapeutic Target Database). Out of the other top candidates we found CHRFAM7A, GRM1, GRM3, ITPR1, HTR7, and COMT particularly interesting: CHRFAM7A is an alpha-7 nicotinic cholinergic receptor subunit interacting with amyloid β, whose aggregates (i.e., plaques) are one of the hallmarks of AD [34]. CHRFAM7A may promote neuronal survival and function, and subunits are expressed by astrocytes participating in synaptic communication [35]. GRM1 is the target of the compound PF-1913539, which has been discontinued in a phase 3 AD trial [31]. GRM3 (mGlu3) is found in astrocytes as well as neuronal cells, and have been observed to have neuroprotective properties. Its agonists and positive allosteric modulators were reported to be potentially helpful for AD treatment [36]. Glial mGlu3 receptors regulate the production of neurotrophic factors such as nerve growth factor, brain-derived neurotrophic factor and glial-derived neurotrophic factor [36]. BCI-632, a compound that targets GRM3, is currently tested in a phase 1 AD trial [31]. ITPR1, an intracellular Ca 2+ channel, mediates calcium release from the endoplasmic reticulum, triggering apoptosis, and its deletion has been linked to spinocerebellar ataxia type 15, a neurodegenerative disease [37, 38]. Single nucleotide polymorphisms (SNPs) rs73310256 in HTR7 [39] and rs4680 in COMT have been associated with AD [40]. COMT is currently discussed as a target for AD [41]. Finally, COMT is the target of the anti-Parkinson drug Tolcapone (source: TTD), supporting our previous finding that GuiltyTargets can re-propose targets from related diseases.

Visualization of interactions between known and candidate targets (Figure 4) revealed that the network of known and proposed targets has a higher than expected interaction rate (PPI enrichment *p* < 1.0*E -* 16, calculated using STRING web interface). Further-more, the candidate targets were observed to reside on the borders of this interaction network. These two observations lead to the hypothesis that targeting the proposed candidates would propagate through the network, influencing disease-related proteins indirectly. Generating drugs that target multiple proteins in this list might be effective, if these candidates were to be considered as different entry points to the disease module.

**Fig 4.**
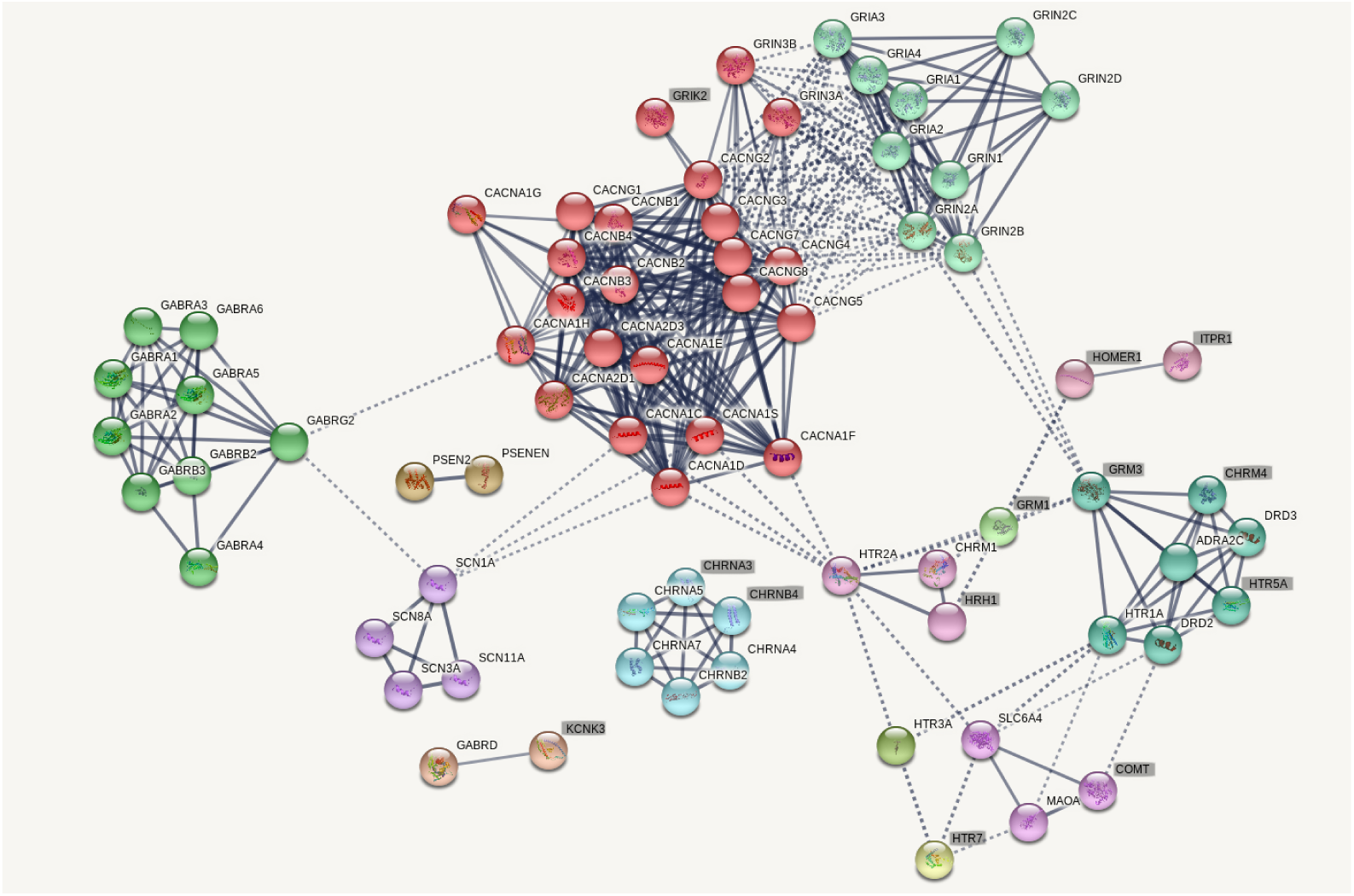
Interactions between known and candidate targets, with confidence scores higher than 0.7. Clusters of nodes were calculated using MCL clustering [42] with inflation parameter of 3.4 and the nodes were colored based on the clusters they are in. The transparency of the links shows the confidence score of the interaction. If a node has some known or predicted 3D structure, it is filled with a structure image. Highlighted nodes show the proposed candidates, whereas the rest show the known targets. Image generated using STRING [29].

## Discussion

We presented a novel network representation learning based approach for target prioritization, GuiltyTargets. Our approach uses a protein-protein interaction network, a differential gene expression profile and a list of known targets to prioritizes proteins as targets for a particular disease. We showed that GuiltyTargets is highly robust and significantly outperforms the method by Emig *et al*. in terms of ranking performance. As demonstrated by our validation studies, it is applicable to various types of diseases, including cancers, metabolic and neurodegenerative diseases. We demonstrated that GuiltyTargets can be used to repurpose existing targets from a different (but related) disease area. Application of GuiltyTargets to AD showed that several of the highest ranked candidates are indeed proposed in the literature for AD, and three of them have been targeted by candidate AD drugs. Moreover, our case study once more demonstrated the possibility to repurpose existing targets from related disease areas with our methods, e.g.COMT as target of the drug Tolcapone in Parkinson’s Disease.

GuiltyTargets as well as other machine learning based target prioritization methods (including the one by Emig *et al*.) learn properties of known targets to rank candidate proteins. Hence, these approaches rely on available information about targets, which is often incomplete and noisy [43]. Because of their dependency on available data machine learning approaches typically have difficulties to propose candidates that point towards completely novel disease biology. Despite these limitations GuiltyTargets showed promising results for target prioritization, including our case study for AD. Hence, we see GuiltyTargets as a promising tool to support the decision process in the context of target identification in pharmaceutical research.

## Materials and Methods

### Information Sources

Three types of information sources were used for target prioritization:

1. differential gene expression profiles between diseased and healthy subjects
2. protein-protein interaction networks (PPI network)
3. disease-specific target annotation

In the following we provide more detailed information about these data.

#### Gene Expression Data

Gene expression data for acute myleoid leukemia, hepatocellular carcinoma, idiopathic pulmonary fibrosis, liver cirrhosis and multiple sclerosis was obtained from Gene Expression Omnibus (GEO) [44], and differential gene expression was assessed via GEO2R [45], Biobase [46], GEOquery [47] and limma [48] using multiple testing correction via the false discovery rate [49] (see: Supplementary Table 1).

For AD, RNASeq data from the AM-PAD Knowledge Portal (AM-PAD) was used [50]. In particular, MSBB, ROSMAP, and MayoRNASeq studies were utilized. Differential gene expression was assessed by applying DESeq2 to the normalized RNAseq data for each brain region (Table S4).

#### Protein-Protein Interaction Networks

As PPI networks, HIPPIE v2.0 [30] and STRING v10.5 [29] were used since both of these networks are created by combining multiple sources of PPIs and provide confidence scores. HIPPIE and STRING differ in the type of interactions they contain (Table S5): HIPPIE relies on physical protein-protein interactions, whereas STRING captures more broadly functional interactions. Hence, STRING has a much larger size than HIPPIE. The analyses on this paper only included the interactions between human proteins. STRING locus identifiers were mapped to Entrez identifies using the mappings provided by STRING.

#### Target Databases

Information about known targets were obtained from two databases: The Therapeutic Target Database (TTD) [31] and Open Targets [18] (see: Table S6). Target identifiers in TTD database were mapped to UniProt identifiers using the conversion file provided by TTD. These identifiers were then mapped to Entrez gene IDs using R packages AnnotationDBI [51] and org.Hs.eg.db [52]. In addition to TTD, known protein targets were retrieved from Open Targets. HGNC symbols were converted to Entrez identifiers using R packages AnnotationDBI [51] and org.Hs.eg.db [52].

### GuiltyTargets

#### Deep Network Representation Learning

GuiltyTargets relies on a deep representation learning of an annotated PPI network via Gat2Vec, where node attributes represent discretized gene expression log fold changes (see Results part). In a first step, two separate graphs, the structural and the attribute graph, are constructed from the original labeled PPI network, where the structural graph corresponds to the PPI network, and the attribute graph is a bipartite graph between protein nodes and discretized log fold changes. Afterwards Gat2Vec retrieves for each vertex its structural context through random walks of a predefined length. The exact parameters that we used to run Gat2Vec are given in Table S7. The result of each random walk is a sequence of vertices and node attributes, respectively. These sequences are subsequently embedded into an Euclidean space using a SkipGram neural network, which is an essential part of the well known Word2Vec method [53].

#### Target Candidate Ranking

The features obtained from annotated PPI networks and the disease-specific target annotations were used to train a logistic regression classifier. Following a PU learning scheme known targets were assigned positive labels, and the remaining proteins were treated as if they were negatives. It is essential to note that proteins that are not known as targets in a particular disease of interest could in fact be targets in another disease context, and finding them is the primary goal target prioritization approaches. For the implementation, the LogisticRegression class from linear model module and OneVsRestClassifier class from multiclass module in Python library scikit-learn were used [54].

## Acknowledgements

We would like to thank Daniel Domingo-Fernández for his valuable help in interpreting the ranked candidate list for Alzheimer’s disease.

The results published here are in part based on data obtained from the AMP-AD Knowledge Portal (doi 10.7303/syn2580853).

For MSBB data set, these data were generated from postmortem brain tissue collected through the Mount Sinai VA Medical Center Brain Bank and were provided by Dr. Eric Schadt from Mount Sinai School of Medicine.

For MayoRNASeq data set, study data were provided by the following sources: The Mayo Clinic Alzheimer’s Disease Genetic Studies, led by Dr. Nilufer Taner and Dr. Steven G. Younkin, Mayo Clinic, Jacksonville, FL using samples from the Mayo Clinic Study of Aging, the Mayo Clinic Alzheimer’s Disease Research Center, and the Mayo Clinic Brain Bank. Data collection was supported through funding by NIA grants P50 AG016574, R01 AG032990, U01 AG046139, R01 AG018023, U01 AG006576, U01 AG006786, R01 AG025711, R01 AG017216, R01 AG003949, NINDS grant R01 NS080820, CurePSP Foundation, and support from Mayo Foundation. Study data includes samples collected through the Sun Health Research Institute Brain and Body Donation Program of Sun City, Arizona. The Brain and Body Donation Program is supported by the National Institute of Neurological Disorders and Stroke (U24 NS072026 National Brain and Tissue Resource for Parkinsons Disease and Related Disorders), the National Institute on Aging (P30 AG19610 Arizona Alzheimer’s Disease Core Center), the Arizona Department of Health Services (contract 211002, Arizona Alzheimer’s Research Center), the Arizona Biomedical Research Commission (contracts 4001, 0011, 05-901 and 1001 to the Arizona Parkinson’s Disease Consortium) and the Michael J. Fox Foundation for Parkinson’s Research.

For ROSMAP data set, study data were provided by the Rush Alzheimer’s Disease Center, Rush University Medical Center, Chicago. Data collection was supported through funding by NIA grants P30AG10161, R01AG15819, R01AG17917, R01AG30146, R01AG36836, U01AG32984, U01AG46152, the Illinois Department of Public Health, and the Translational Genomics Research Institute.

## Author contributions statement

Ö.M. and C.T.H. implemented the method and analysed data. M.H.-A. and H.F. designed the research. H.F. supervised the project. OÖ.M., C.T.H. and H.F. drafted the manuscript.

